# The Sync/deSync model: How a synchronized hippocampus and a de-synchronized neocortex code memories

**DOI:** 10.1101/185231

**Authors:** George Parish, Simon Hanslmayr, Howard Bowman

## Abstract

Neural oscillations are important for memory formation in the brain. The de-synchronisation of Alpha (10Hz) oscillations in the neo-cortex has been shown to predict successful memory encoding and retrieval. However, when engaging in learning, it has been found that the hippocampus synchronises in Theta (4Hz) oscillations, and that learning is dependent on the phase of Theta. This inconsistency as to whether synchronisation is ‘good’ for memory formation leads to confusion over which oscillations we should expect to see and where during learning paradigm experiments. This paper seeks to respond to this inconsistency by presenting a neural network model of how a well-functioning learning system could exhibit both of these phenomena, i.e. desynchronization of alpha and synchronisation of theta during successful memory encoding.

We present a spiking neural network (the Sync/deSync model) of the neo-cortical and hippocampal system. The simulated hippocampus learns through an adapted spike-time dependent plasticity rule, in which weight change is modulated by the phase of an extrinsically generated theta oscillation. Additionally, a global passive weight decay is incorporated, which is also modulated by theta phase. In this way, the Sync/deSync model exhibits theta phase-dependent long-term potentiation and long-term depression. We simulated a learning paradigm experiment and compared the oscillatory dynamics of our model with those observed in single-cell and scalp-EEG studies of the medial temporal lobe. Our Sync/deSync model suggests that both the de-synchronisation of neo-cortical Alpha and the synchronisation of hippocampal Theta are necessary for successful memory encoding and retrieval.

**Significance statement:** A fundamental question is the role of rhythmic activation of neurons, i.e. how their firing oscillates between high and low rates. A particularly important question is how oscillatory dynamics between the neo-cortex and hippocampus contribute to memory formation. We present a spiking neural-network model of such memory formation, with the central ideas that 1) in neo-cortex, neurons need to break-out of an alpha oscillation in order to represent a stimulus (i.e. alpha desynchronises), while 2) in hippocampus, the firing of neurons at theta facilitates formation of memories (i.e. theta synchronises). Accordingly, successful memory formation is marked by reduced neo-cortical alpha and increased hippocampal theta. This pattern has been observed experimentally and gives our model its name – the Synch/deSynch model.

## Introduction

Brain oscillations, via their ability to synchronize and desynchronize neuronal populations, play a crucial role in the formation and retrieval of episodic memories. However, little is known about how oscillations implement the necessary mechanisms for encoding and retrieval of such memories. This knowledge gap is partly due to a lack of computational models simulating oscillatory behaviours as observed in human EEG/MEG recordings during memory tasks. The link between oscillations and memory is further complicated by empirical data, which has fuelled a conundrum as to how oscillations relate to memory. Specifically, hippocampal theta (^~^3-8 Hz) and gamma (^~^40-80 Hz) synchronisation (Fell & Axmacher, 2011) and the de-synchronisation of alpha and beta (8-30 Hz) in cortical regions (Hanslmayr, et al., 2012) have both been reported as important for memory encoding and retrieval. Classic computational models theorise that hippocampal and neo-cortical regions offer functionally distinct mechanisms to form episodic memory (O’Reilly, et al., 2014), where a sparsely connected hippocampus learns new information quickly and a dense neo-cortex incorporates this information slowly. Building on these complementary learning systems we recently presented a potential solution to the synchronization/de-synchronization conundrum (Hanslmayr, et al., 2016), suggesting that hippocampal theta synchronisation (^~^4Hz) mediates the binding of concepts, while neocortical alpha de-synchronisation (^~^10Hz) is due to the representations of these concepts becoming active. We here present a first computational network model which implements these mechanisms and simulates the opposing synchronizing and desynchronizing behaviours in the hippocampus and neocortex during a typical episodic memory task. Our model, while being very simple, successfully simulates a number of empirical findings ranging from human single neuron recordings, intracranial EEG recordings, to non-invasive EEG/MEG recordings and therefore represents a useful theoretical link between different levels of human electrophysiological recordings.

Theta oscillations in medial temporal lobe are assumed to play a key role in the formation of memories, where learning is dependent on the power of theta oscillations and the timing of action potentials in relation to the ongoing theta cycle (Rutishauser, et al., 2010) (Backus, et al., 2016) (Staudigl & Hanslmayr, 2013) (Heusser, et al., 2016). Studies in rodents have provided a mechanism by which theta oscillations exert their influence on memory in showing that Long-Term-Potentiation (LTP) and Long-Term-Depression (LTD) occur at opposite phases of the theta cycle (Huerta & Lisman, 1995) (Pavlides, et al., 1988). Building on theories of synaptic plasticity, it has been postulated that LTD occurs whilst most neurons in region CA1/CA3 are active in the excitatory phase of theta (as recorded from CA1/CA3 hippocampal regions), whereas LTP occurs in the inhibitory phase of theta when most neurons are silent (Hasselmo, 2005). We henceforth refer to maximum excitation and inhibition during these phases of theta as the peak and trough, respectively. The model we describe here shows that stimulated hippocampal cells demonstrate a phase shift forward in theta, enabling LTP to occur in the trough of theta where other non-stimulated cells are silent.

Concerning alpha oscillations, it can be assumed that there is a negative relationship between alpha power and discriminating neural activity (Haegens, et al., 2011), leading to the notion that alpha provides functional inhibition (Klimesch, et al., 2007) (Jensen & Mazaheri, 2010). Supporting this notion, alpha power decreases (i.e. desynchronizes) are often localized in cortical regions relevant for a given task, whereas alpha power increases occur in competing areas which are being inhibited (Jokisch & Jensen, 2007) (Wardhauser, et al., 2012). These findings suggest that the de-synchronisation of alpha represents the flow of information to a targeted group of neurons. Consistent with this general gating function of alpha, power decreases are strongly evident in episodic memory tasks where cortical alpha power decreases predict successful encoding (Hanslmayr, et al., 2012) and retrieval (Khader, et al., 2010) (Waldhauser, et al., 2016). In addition to the hippocampal theta dynamics, our model also simulates the memory dependent alpha power decreases in the neocortex during the encoding and retrieval of episodic memories.

## Materials and methods

### Computational model

Here we describe a simple computational neural network model, which takes inspiration from the complementary learning systems framework (CLS), and lends credence to the previously theorised notion that opposing oscillatory behaviour in cortical and Hippocampal regions both contribute to episodic memory formation (Hanslmayr, et al., 2016). We do not describe in detail complex Hippocampal, entorhinal or neo-cortical cell assemblies, as in other models (Hasselmo & Eichenbaum, 2005) (O’Reilly, et al., 2014), choosing instead to model a simple mechanism to simulate a typical episodic memory paradigm where an association between stimuli has to be learnt in one trial. A principal of our modelling endeavour has been to identify the simplest neural instantiation of our theory under an Ockham’s razor principle.

### Experimental paradigm

We chose to compare our model to an experiment that recorded from medial-temporal-lobe (MTL) neurons within epilepsy patients (Ison, et al., 2015). As depicted in Figure 1A, the experimenters screened many images of people to each participant for the purpose of finding one that the neuron under observation responded to, denoted from here on as the preferred (P) image. A separate image of a location was chosen that the neuron did not respond to, denoted as the non-preferred (NP) image. The P image of the person was then digitally superimposed onto the NP image of the location (denoted here as the composite (C) image), before being presented to the participant in what is termed here as the learning phase. The experimenters then conducted the screening process again, presenting both the NP & P images, to assess the impact of learning on the activity of the Hippocampal neuron. Figure 1A shows how we simulated this paradigm, where there is a screening phase before and after the presentation of the composite stimulus.

**Figure 1:**
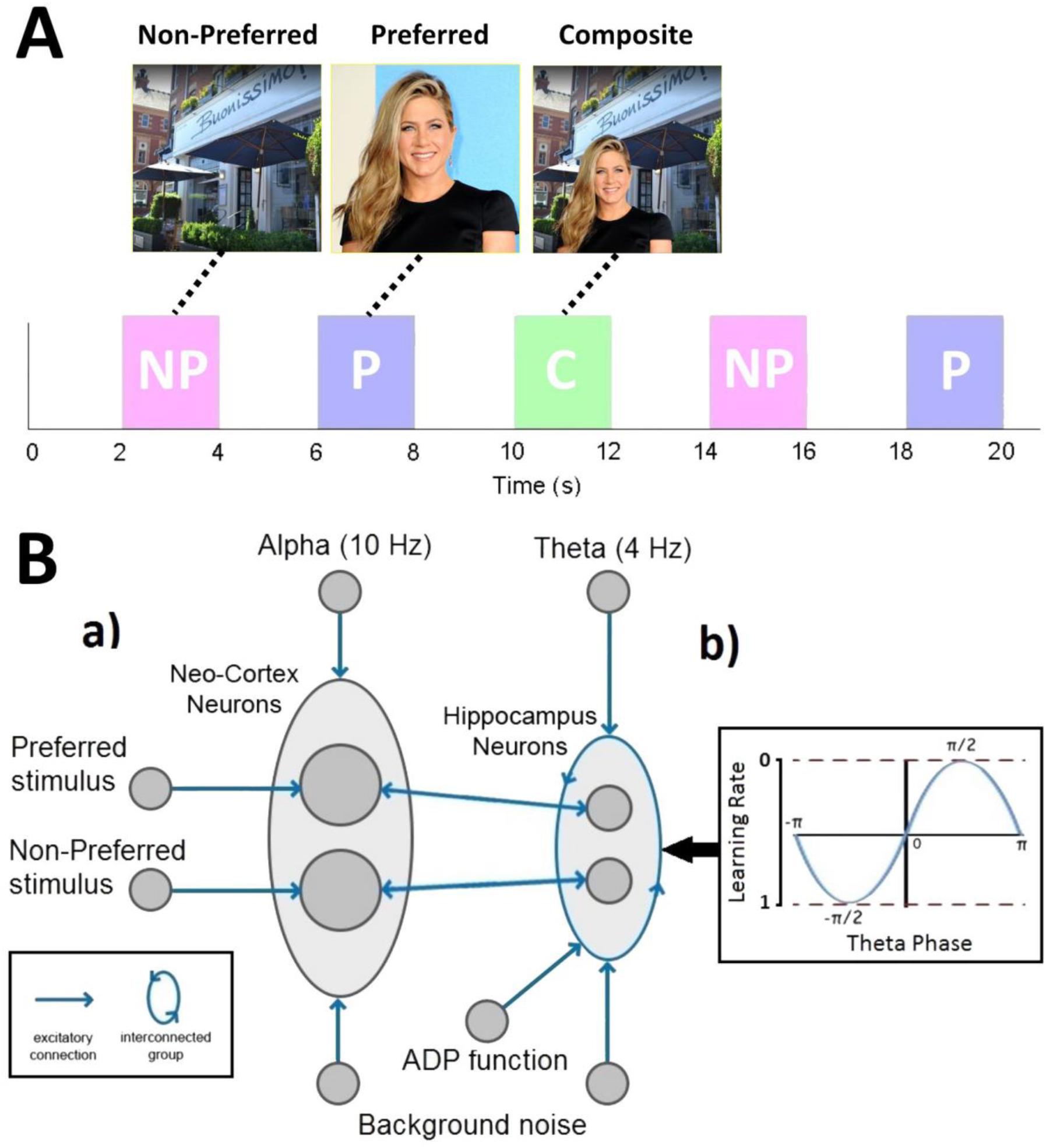
Experimental paradigm **(A)**. A non-preferred (NP) and preferred (P) image are found that the neuron does not and does respond to. These are then combined and presented in a composite (C) stimulus. Both P and NP images are presented again after this learning phase. Network connectivity **(B)**. The architecture of the network **(a)** shows how a group of neo-cortical (NC) neurons and a group of Hippocampal neurons receive input from a 10Hz and 4Hz tonic wave, respectively, and both groups receive (background) noise from a Poisson distribution of spikes. Two subgroups of NC neurons receive input from higher level areas that represent the P and NP image. Each subgroup of NC and Hip neurons have reciprocal connectivity between themselves. Hippocampal neurons also receive an after-de-polarisation (ADP) function. Hippocampal neurons are interconnected, and spike-time-dependent-plasticity (STDP) is enabled with a Theta phase dependent learning rate **(b)**.

### Neuron physiology

Our model comprises two groups of neurons representing the neo-cortex (NC) and the hippocampus (Figure 1Ba), split again into two subgroups coding for the P and NP images (where the number of neurons in each group was *N*_*NC*_ = 20, *N*_*hip*_ = 10). All neurons are simulated using an Integrate-and-Fire equation (*equations* 1, *V*_*th*_ = −55*mV*, *E* = −70*mV*, *C*_*m*_ = 240*nF*, *V*_*ref*_ = 2*ms*, *τ*_*m*_ = 20*ms*). A spike event is sent to other downstream connected neurons if the membrane potential (*V*_*m*_(*t*)) of a neuron surpasses the threshold for firing (*V*_*th*_). After a spike, the neuron goes into a refractory period, where the membrane potential is clamped to the resting potential (*E*) for a set period (*V*_*ref*_). With this equation, the membrane potential of a neuron is constantly decaying to its resting potential (*E*) at a rate dictated by the membrane time constant (*τ*_*m*_). The sum of all inputs at *t* is divided by the capacitance (*C*_*m*_) of the membrane potential. Inputs originate from constant alternating currents (*I*_*tonic*_), the sum of excitatory-post-synaptic-potentials (EPSPs) from spikes at each input synapse (*I*_*syn*_) and an after-de-polarisation function (*I*_*ADP*_), which will be described in more detail later.

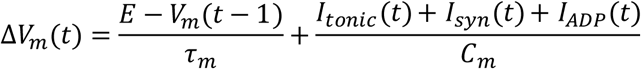

Equation 1: The integrate-and-fire model

An alpha function (*equation* 2) was used to model EPSPs for incoming spike events, where Δ*t* is equal to the current time (*t*) minus the time of the eliciting spike (*t*_*fire*_). The higher the synaptic time constant *τ*_*s*_, the larger the integral of the EPSP, ensuring that a spike has a more sustained effect on the receiving neuron’s membrane potential. As shown in Figure 1Ba, bi-directional excitatory connections coupled each P & NP subgroup between the NC and Hippocampus. Synapses from the NC had a smaller constant (*τ*_*s*_ = 1.5*ms*) with a larger weight, whilst synapses from the Hippocampus had a larger constant *τ*_*s*_ = 15*ms*) with a smaller weight. Therefore, spikes from the NC to the Hippocampus had an immediate and large effect, whilst spikes in the opposite direction had a smaller yet more sustained effect, emphasising their ability to cause ERD of alpha frequencies upon memory retrieval. Synapses within the Hippocampus also had slightly larger synaptic time constants (*τ*_*s*_ = 5*ms*) to allow them to more easily interact with one another. Spikes originating from external noise generators had a synaptic time constant of 1.5*ms*.

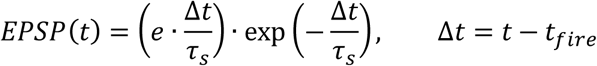

Equation 2: The Excitatory-Post-Synaptic-Potential (EPSP)

### Neocortical system

Based on CLS, the NC system learns slowly from repeated presentations. As our model emphasises the effect of oscillations on a single learning event, we assumed the existence of two pre-established NC populations, one representing the P and the other the NP concept, where synaptic modification was not implemented due to an assumed slow cortical learning rate, and there was no connectivity between neurons in each population (Figure 1Ba). Each NC neuron received background noise, representing “chatter” from other brain regions, in the form of a Poisson distributed spike-events (^~^45k spikes/s). We do not explicitly model a neural mechanism for oscillations, thus a cosine wave of frequency 10Hz (amplitude = 21pA) was fed into NC neurons via *I*_*tonic*_ to model ongoing Alpha. Two separately generated Poisson distributed spike-trains (^~^100k spikes/s) were then paired with each NC subgroup upon stimulus presentation, modelling the activation of the P and/or NP images from higher cortical and visual areas.

### Hippocampal system

Hippocampal neurons were similarly organised into two subgroups (Figure 1Ba), where each neuron received background noise (4k spikes/s) and a cosine wave of 4Hz (amplitude = 25pA) to model ongoing Theta. Based on CLS, the Hippocampal system learns quickly from a single presentation. Therefore, Hippocampal synaptic modification was enabled via an adapted Spike-Time-Dependent-Plasticity (STDP) learning rule (Song, et al., 2000). We adjusted this rule to relate to empirical evidence that Hippocampal learning is Theta phase dependent (Huerta & Lisman, 1995), with LTD occurring in the peak and LTP in the trough of theta, as recorded in CA1/CA3 (Hasselmo, 2005). To this end, synaptic LTP was implemented by multiplying STDP weight modifications by the phase of the Theta cosine wave, with a value between 0 and 1 during the peak and trough, respectively (Figure 1Bb).

When a neuron spiked, a reward (*A*_+_) for contributing synapses was calculated as the product of a constant learning rate (*ε* ∈ ℝ. 0 ≤ *ε* ≤ 1), Theta at time *t* (*θ* ∈ ℝ. 0 ≤ *θ* ≤ 1) and the maximum weight (*W*_*max*_), whilst punishments for competing synapses were calculated as *A*_ − = 1.1 · *A*_+_ (*equation* 3). The greater strength for *A*_ compared to *A*_+_ reflected a preference for synaptic weakening in order to maintain a stable network. Whenever a spike event occurs, at unit *i* or *j*, an accumulated STDP update *v*_*ij*_(*t*) for synapse *i* to *j* is calculated from its history of previous spiking (*i* then *j* or *j* then *i*) (*equation* 5). A function was then used to calculate the STDP acting on the synapse (*equation* 4), where an exponential weighting of *A*_+_ was applied if the pre-synaptic spike occurred before the post-synaptic spike and of *A*_ if the post-synaptic spike occurred first. All Hippocampal weights were subject to STDP updates, along with an exponential passive decay, which was multiplied by the negative phase of Theta (1 ‒ θ(t)) (*equation* 6). The presence of this decay is consistent with the non-specific LTD that might occur during oscillatory spiking in the peak of Theta (Hasselmo, 2005). This decay was larger for smaller weights, establishing a transition point whereby weakly interacting synapses were pruned (*τ*_*w*_ = 20). A piecewise linear bounding function was also used to protect against sign reversal and run-away weights (*equation* 7; *W*_*max*_ = 120; *W*_*min*_ = 0).

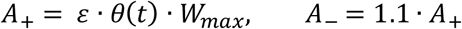

Equation 3: Reward (*A*_+_) and punishment (*A*_) of synapses.

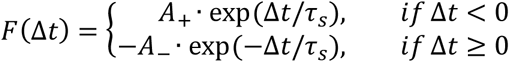

Equation 4: Function for STDP between pre and post-synaptic spikes (Song, et al., 2000), where Δ*t* is always the difference between the time of a pre-synaptic and post-synaptic spike.

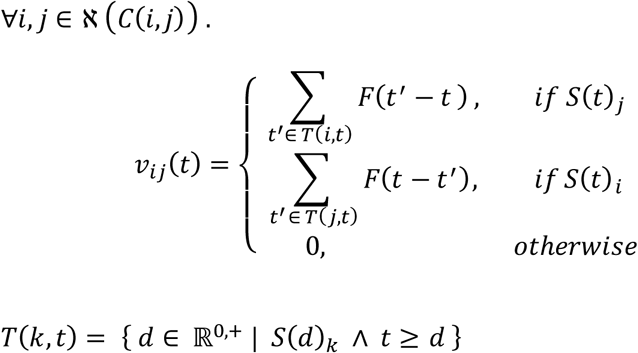

Equation 5: SDTP synaptic modification at time *t* for a network with node labels ℵ = {1,…,*n*}. *C*(*i,j*) is true if and only if *i* and *j* are connected. *S*(*t*)_*i*_ indicates a spike event at the *i*th neuron at time *t.T*(*k*) determines the set of all times, at which there was a spike at neuron *k* at a time before *t*. This is used to provide spike events paired, across synapse *i*, *j*, with the spike at time *t*. In addition, we use auxiliary weight variables *v*_*ij*_ and *V*_*ij*_ to enable application of a piecewise linear bounding function, see eqn 7.

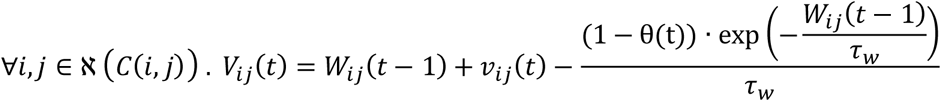

Equation 6: Update of auxiliary weight variable and implementation of non-specific passive decay of synapses.

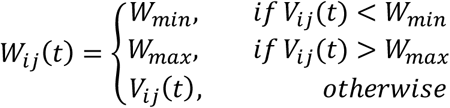

Equation 7: Piecewise linear bounding function

Hippocampal neurons were interconnected with a probability of 40% to form a connection. Additionally, as it was assumed that both images were previously known to the participants but not associated, a random 50% of synapses within each subgroup had initial synaptic weights of *W*_*max*_ whilst all others were set to 0. This ensured the random assignment of pre-established sets of winning and losing pathways within the subgroups coding for the P & NP image.

Hippocampal neurons received additional input from an After-De-Polarisation (ADP) function (Jensen, et al., 1996) to control activation (*equation* 8; *A*_*ADP*_ = 100*pA*, *τ*_*ADP*_ = 250*ms*). This provided exponentially ramping input, which was reset after each spike-event (*t*_*fire*_). Evidence for an ADP function in hippocampal neurons has been found experimentally during cholinergic (Andrade, 1991) (Caesar, et al., 1993) (Libri, et al., 1994) and serotonergic (Araneda & Andrade, 1991) modulation, and has the effect here of modelling an effectively inhibitory input for each Hippocampal neuron, which wanes the further one is from the eliciting spike.

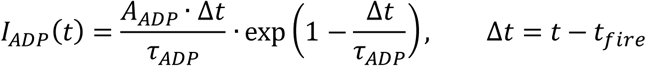

Equation 8: After-De-Polarisation (ADP) function

### Local Field Potential (LFP) and Time Frequency Analysis (TFA) methods

The LFP measures the activity of a group of neurons by first aggregating spikes through time. This was then filtered twice, first by using a Hanning filter with a 30ms window and then again with a sampling frequency between 2-6Hz or 8-12Hz dependent on whether we are filtering by Theta or Alpha, respectively. The LFP was analysed in time-frequency space using a Gabor filter with an upper and lower bound of 2-6Hz or 8-12Hz for Theta or Alpha analysis (γ = 0.5 for <30Hz or γ = π/2 for >30Hz). The absolute values were then taken and plotted in time-frequency space.

## Results

### Simulation procedure

We simulated our model based on a learning paradigm used in an MTL single cell recording experiment (Ison, et al., 2015). During the initial screening phase, both the P & NP images were presented individually. This was simulated by independently creating two Poisson distributed spike trains (^~^100k/s for 2 seconds) that fed into each respective P & NP subgroup of NC neurons (Figure 1A; P = blue, NP = magenta). An inter-stimulus interval of 2 seconds was used. Afterwards, we presented both images in a composite stimulus (green), where both subgroups of NC neurons concurrently received spike-trains. Following this learning phase, we repeated the screening phase to assess the capability of the network to associate these stimuli together. The whole process was simulated 1000 times to assess the variability of the network, where for each simulation the Alpha and Theta cosine waves began at a different random phase and new noisy spike trains were generated. The following results take an average over all simulations.

### Hippocampal weight change

Maximal synaptic modification occurs between Hippocampal neurons that are stimulated to shift forward in phase and fire towards the trough of an ongoing Theta oscillation (Hasselmo, 2005). Due to this, synaptic modification only occurs during the screening and learning phases of the stimulation (Figure 2; NP stimulus-magenta; P stimulus-blue; C stimulus-green) and not during the inter-stimulus intervals. Due to the maximisation of a random 50% of synapses within each P & NP subgroup, the average weights of these groups begin at *W*_*max*_/2 (Figure 2A). Throughout the entire simulation, there is weight change within each subgroup (P-blue line; NP-magenta dash) when the respective image they are coding for is presented. With the competitive STDP rule, winning and losing weights are pushed towards *W*_*max*_ or *W*_*min*_ respectively, causing a capping effect where a weight in one direction can still change whilst its competitor is capped. Here, this means that the average weight of each subgroup rises a small amount to stabilise just above *W*_*max*_/2 every time the respective image is presented, effectively reinforcing both P & NP concepts.

**Figure 2:**
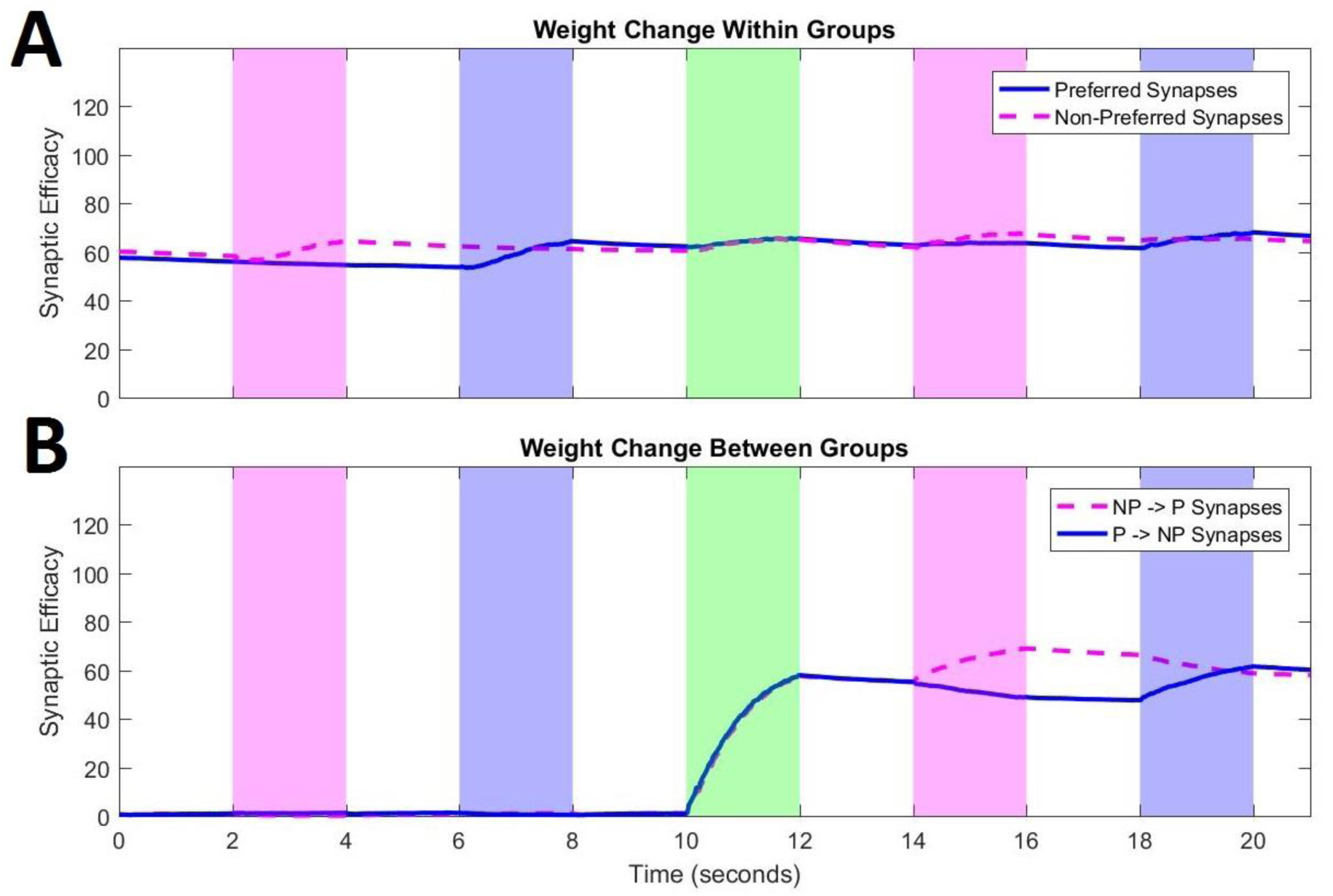
Hippocampal weight change throughout the simulation both within **(A)** and between subgroups **(B)** that code for the P and NP stimulus. Weights within each subgroup increase when the relevant image is presented **(A)**, where the magenta and blue periods indicate the presentation of the NP and P images, respectively, and the green period indicates the presentation of both images that have been combined into a composite image. During this learning period, weights from the NP to the P subgroup (magenta dashed) and vice-versa (blue solid) increase **(B)**. Outgoing weights then increase upon the presentation of the relevant stimulus.

When the composite stimulus is presented (green), there is only marked synaptic change between both subgroups (Figure 2B; P->NP-blue line; NP->P-magenta dash). Here, weights go up bi-directionally as both subgroups of neurons are concurrently stimulated to become active towards the trough of Theta. When the screening phase is repeated after the learning phase, weights increase in the direction from the active population to the non-active population and decrease in the reverse direction. This is due to STDP competition taking place within a pool of shared synapses, where winning pathways that once spanned across neurons in both directions during the learning phase, are pulled in a direction whereby the most active neurons are presynaptic during the 2^nd^ screening phase.

Weights passively decay very slowly according to an exponential pattern to model the effect of a large population of neurons spiking during the peak of Theta, where LTD has been found to occur (Hasselmo, 2005). As LTP occurs over a spectrum of 1 to 0, small weight increases occur as neurons spike on either side of the peak of Theta. The passive decay implemented here is stronger for smaller weights (equation 6), to mitigate these gradual weight increases and prune irrelevant synapses. LTD weight decay is also prominent in the inter-stimulus periods, where all weights slowly reduce over time.

### Hippocampal activity

Activity is measured as the average output of spikes per subgroup (10 neurons per item) that occur within a time period. Figure 3 shows activity of the Hippocampal preferred (P) subgroup at different stages of the simulation, taking an average over 1000 simulations to mitigate the random phase starts of Theta and Alpha oscillations, using a bin width of 20ms. The mean firing rate is shown with bootstrapped confidence intervals.

**Figure 3:**
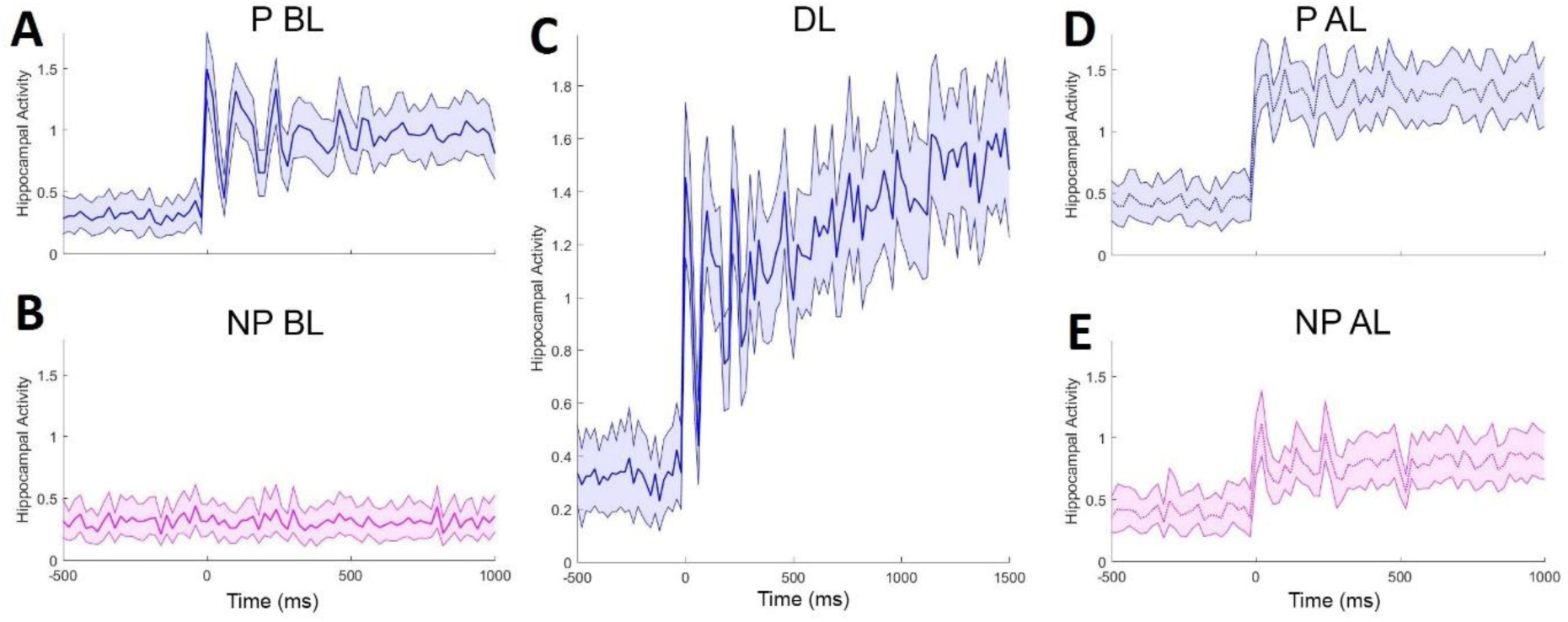
Activity of the Hippocampal subgroup encoding for the preferred (P) image, when both the P **(A)** and NP **(B)** images are presented before the learning phase **(C)**, and then the P **(D)** and NP **(E)** images are presented again afterwards. Increasing weight change during the learning phase **(C)** causes gradually increasing activity. This increase in weights between both subgroups then causes the P subgroup to respond to the NP image **(E)**.

During the first screening phase, P neurons become active when the P image is presented but not when the NP image is presented (Figure 3A-B; P before learning (BL) – blue, NP before learning (BL)-magenta). Stimulus onset phase resets the ongoing Theta oscillation, which then synchronises the ADP function, generating a damped oscillation that is phase consistent across replications (Figure 3A, 0-500ms). After a few cycles, activity averages out across replications, as neurons re-synchronise ADP input with their relative 4Hz tonic input.

A similar pattern occurs when the C image is presented during learning (Figure 3C), where not only is there an oscillatory evoked transient but activity increases over time. This is due to the large amount of weight change occurring between the P & NP subgroups (Figure 2B; 10-12s), with activity feeding back into each subgroup.

After this learning phase, the P & NP images are independently presented again (Figure 3D-E; P after learning (AL) – blue dashed, NP after learning (AL), magenta dashed). Due to the aforementioned weight change, the P subgroup now responds when the NP image is presented, replicating experimental evidence from a recent human single unit learning paradigm (Ison, et al., 2015). This result also shows that our model efficiently learns associations between two arbitrary stimuli in one short presentation, a crucial requirement for a model of episodic memory. It is interesting to note the slight increase in activity pre-stimulus (Figure 3D-E; -500-0ms), caused by increased weights between the P & NP subgroups leading to feedback activation. This is followed by the typical burst of activity at stimulus onset (0-50ms), yet the aforementioned damped oscillation is mitigated due to this reciprocal feedback of activity between P & NP subgroups negating the ADP function’s recovery period (50-100ms). In experimental research, encoding neurons become less active with successive presentations of the same stimulus (Ison, et al., 2015), perhaps due to a repetition suppression mechanism (Pedreira, et al., 2010). In our model, the P subgroup responds more strongly to the P image when it is presented again after the learning phase, due to an increase in weights causing feedback activation. This could be countered by implementing a habituation mechanism that lies outside of the scope of this model.

### Theta phase

Figure 4 shows Theta phase within the P subgroup of Hippocampal neurons, in response to the presentation of the NP image during the 3 stages of the simulation. The red and green halves of the polar distribution represent the excitatory and inhibitory phases of the 4 Hz cosine wave used to model Theta, where 0° is the peak and 180° is the trough of Theta as recorded in the Hippocampal region CA1/CA3 (Hasselmo, 2005). The total number of spikes occurring within each phase quadrant of Theta was recorded (Figure 4Aa-c), as well as the first spike of each neuron after maximum inhibition (>180°) (Figure 4Ba-c). The latter analysis was performed to show how Hippocampal neurons shift forward in Theta phase once stimulated. Spike numbers were normalised over 1000 simulations.

**Figure 4:**
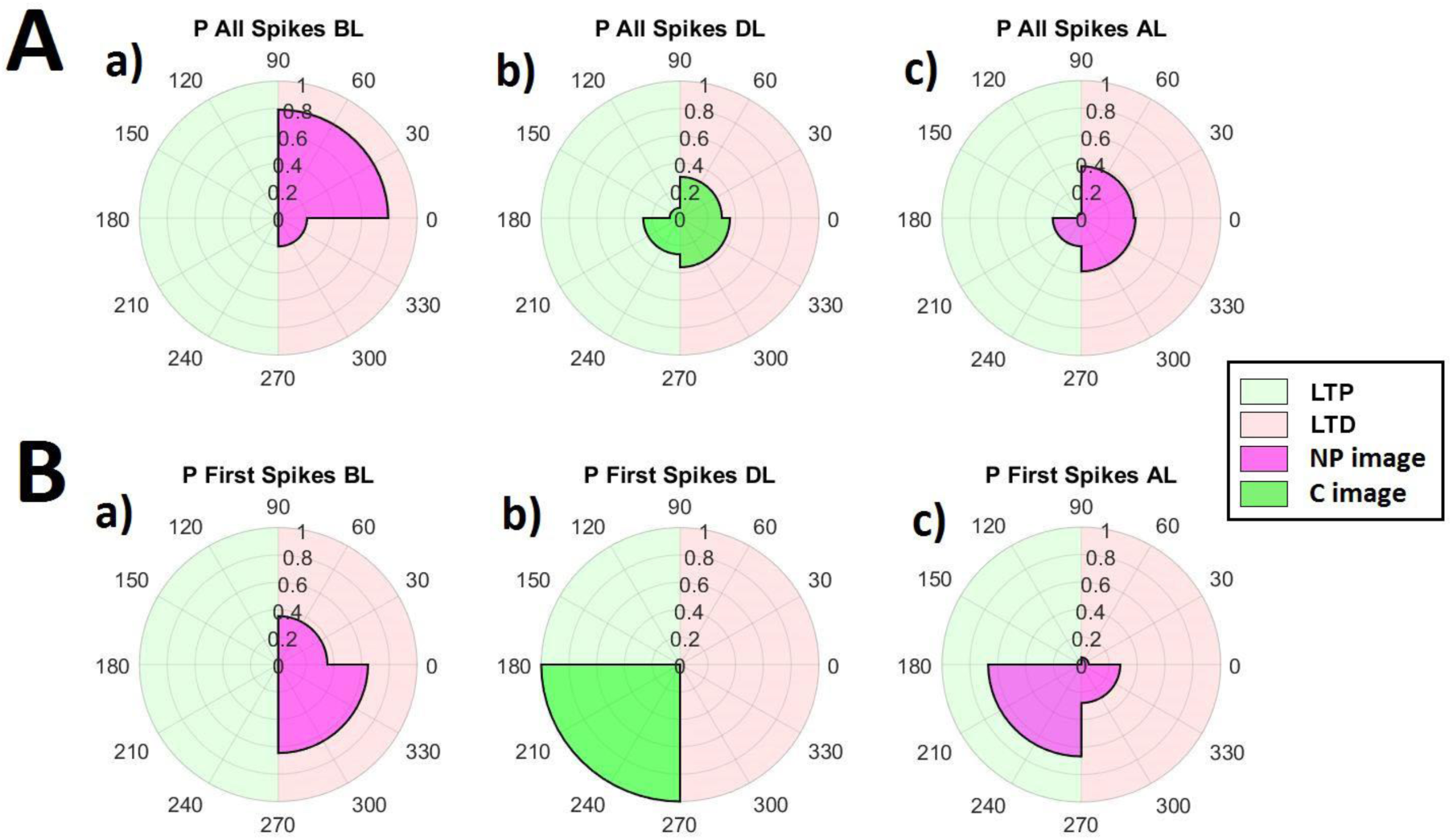
Polar histogram of all spikes **(A)** from the preferred (P) Hippocampal subgroup before learning (**a** BL), during learning (**b** DL) and after learning (**c** AL). Theta is maximal and minimal at 0 and 180 degrees, where LTD and LTP occur in the red and green halves, respectively. The same analysis is shown for the first spikes of each neuron during a Theta phase **(B)**.

Before learning, P neurons are un-responsive to the NP image and oscillate at Theta, where most spikes occur just after maximum excitation (Figure 4Aa; 0-90°), with the first spikes generally occurring just before maximum excitation (Figure 4Ba). When the C image is presented during the learning phase, the P subgroup becomes active across all phases of Theta (Figure 4Ab). Importantly, in order for activation to overcome inhibition, more activity will occur during the peak of Theta. The least number of spikes occur early on in the inhibitory phase (90-180°) due to the effect of the ADP function, which prolongs the recovery time of neurons that spiked late during the excitatory phase (0-90°). Neurons also exclusively spiked first immediately after the trough (Figure 4Bb; 180-270°), indicating that all neurons in the P subgroup successfully phase-shifted forward once stimulated during learning.

When the NP image is presented again after learning, P neurons now respond. Spikes occur in most phase quadrants of Theta (Figure 4Ac), mostly during the excitatory phase as inhibition is overcome, allowing spikes to first occur during the negative phase of Theta (Figure 4Bc) and demonstrating a phase shift forward in Theta. This shift in phase is an index of successful learning and has been well documented in rodents for neurons encoding a particular place when the rodent approaches that place (Huxter, et al., 2003). Our model shows a similar behaviour and predicts that this shift in phase is responsible for associative memory formation. Importantly, this phase shift is most evident when analysing only the first spike within a theta cycle, starting at the theta trough (i.e. where inhibition is maximal). This prediction can be tested in studies recording single units and local field potentials in human epilepsy patients (Ison, et al., 2015).

### Alpha De-Synchronisation

Figure 5Aa-e shows time-frequency power spectra (8-12Hz) of the LFP of the NC P subgroup of neurons throughout each stage of the simulation. A thick band at 10Hz shows non-stimulated NC P neurons oscillating at Alpha (Figure 5Aa), as they do not respond to the NP image at this time. When the P image is presented, a de-synchronisation of Alpha is exhibited (Figure 5Ab/c/e; 0-1s), simulating the well-documented effect of Alpha suppression upon visual stimulation (Berger, 1929). The same effect can be seen when the C image is presented, replicating findings that Alpha de-synchronisation can predict successful memory encoding (Hanslmayr, et al., 2012). A similar, but weaker effect can be seen when the NP image is presented after learning (Figure 5Ad; 0-1s). This de-synchronisation is due to learning driven activation of Hippocampal neurons caused by the association to the NP stimulus. This low-frequency drive (from Hippocampus to Neo-cortex) is magnified by the high time constant on those synapses (*τ*_*s*_), thus de-synchronising Alpha by causing substantial activation in the inhibitory phase. The effect can be more clearly seen in Figure 5B, where a 10% relative decrease in Alpha power from pre to post stimulus is exhibited (Figure 5Bb; 0-1s), consistent with the findings that memory retrieval can be predicted by this same Alpha de-synchronisation (Hanslmayr, et al., 2012). Pre stimulus Alpha power is also slightly weaker (Figure 5Aa; -1-0s), due to stronger weights within the P Hippocampal subgroup causing more activation.

**Figure 5:**
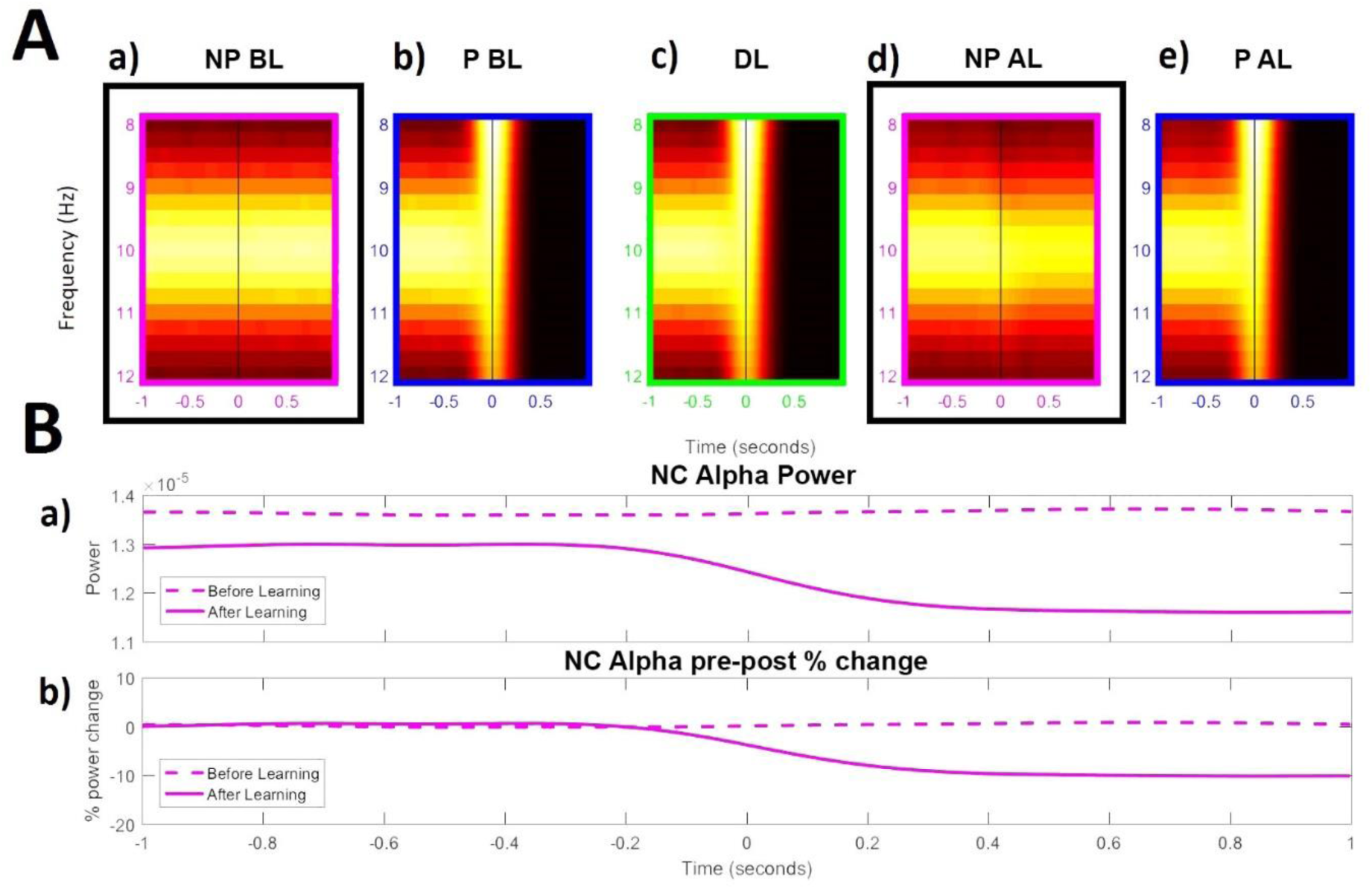
Time-frequency-analysis (TFA) of Neo-Cortical Alpha for the preferred (P) subgroup **(A)** when the non-preferred (NP) **(a)** and P **(b)** images are presented before learning (BL), during learning (DL) **(c)**, and both presented individually again **(d-e)** after learning (AL). A time-course of Alpha power is shown **(B)** for the highlighted boxes **(Aa, Ad)**, where pure power **(Ba)** and percent change in pre-post stimulus power **(Bb)** are shown.

This behaviour of our model mimics several findings in the literature showing memory dependent Alpha power decreases during the reinstatement of episodic memories (Khader, et al., 2010) (Waldhauser, et al., 2016) (Michelmann, et al., 2016). Here, the de-synchronisation of Alpha represents the flow of information in the NC caused by activation of relevant stimuli (Jensen & Mazaheri, 2010), (Klimesch, et al., 2007).

### Theta Synchronisation

Figure 6Aa-e shows time-frequency power spectra (2-4Hz) of the LFP of the Hippocampal P subgroup of neurons throughout each stage of the simulation. Initially, P neurons do not respond to the NP image and oscillate at Theta (Figure 6Aa). An increase in Theta power accompanies increased activation, as P neurons respond to the P image before and during learning (Figure 6Ab-c). Theta synchronisation is stronger during learning, corresponding with experimental evidence (Backus, et al., 2016) (Lega, et al., 2012) (Staudigl & Hanslmayr, 2013). This is due to the rapid increase in synaptic weights during this period (Figure 2B; 10-12s) causing feedback activation, which, in turn, causes more neurons to fire above threshold, but according to the theta rhythm.

**Figure 6:**
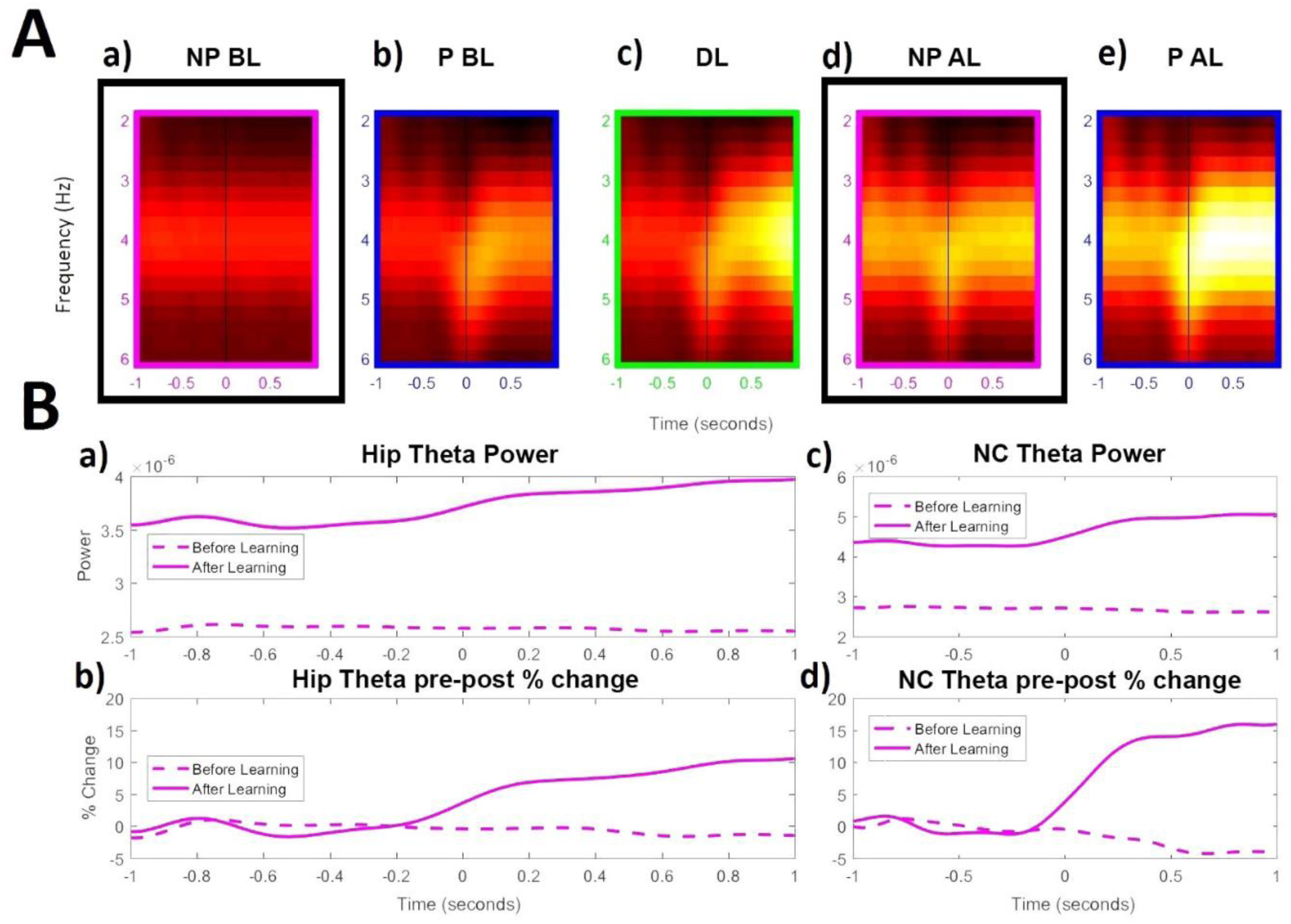
Time-frequency-analysis (TFA) of Hippocampal Theta for the preferred (P) subgroup **(A)** when the non-preferred (NP) **(a)** and P **(b)** images are presented before learning (BL), during learning (DL) **(c)**, and both presented individually again **(d-e)** after learning (AL). A time-course of Theta power is shown **(B)** for the highlighted boxes **(Aa, Ad)**, where pure power **(Ba)** and percent change in pre-post stimulus power **(Bb)** are shown. The same analysis is shown for neo-cortical Theta power during the same time periods **(Bc-d)**.

After the learning phase, P neurons are responsive to the NP image, where a synchronisation of Theta occurs due to an increase in activity post stimulus (Figure 6Ad). This can be seen more clearly in Figure 6Bb, where there is up to a 10% increase in Theta power relative to the pre stimulus period. Due to stronger weights between the P & NP cluster, there is increased feedback activity during the normal oscillatory rhythm. This activity is amplified by a higher synaptic time constant, causing an increase in pre stimulus Theta power (Figure 6Ba; -1-0s). The same changes in Theta power are passed through to the NC (Figure 6Bc-d), which is consistent with experimental evidence of increases of Theta in NC areas after learning paradigm experiments (Burke, et al., 2014) (Klimesch, et al., 2005).

### Varying Stimulus Strength

We next varied how strongly our simulated participant perceived the P & NP images during the screening phases, allowing us to explore the sync/de-sync of Hippocampal Theta and NC Alpha over time at different strengths. This is achieved by varying stimulus strength, i.e. the rate of spikes per second being fed into NC neurons at stimulus onset, and taking the average power during the post-stimulus period across frequencies (0-30Hz). It can be shown that for weakly perceived stimuli, the NC actually synchronises in Alpha within the model (Figure 7A). This is due to input activity being too weak to overcome the trough of the 10Hz cosine input, but strong enough to cause more spiking in the peak. As stimulus strength increases, a de-synchronisation of Alpha is obtained as neurons overcome inhibition to spike across all phases of Alpha, indicating why Alpha de-synchronisation is a marker of successful memory encoding (Hanslmayr, et al., 2012).

**Figure 7:**
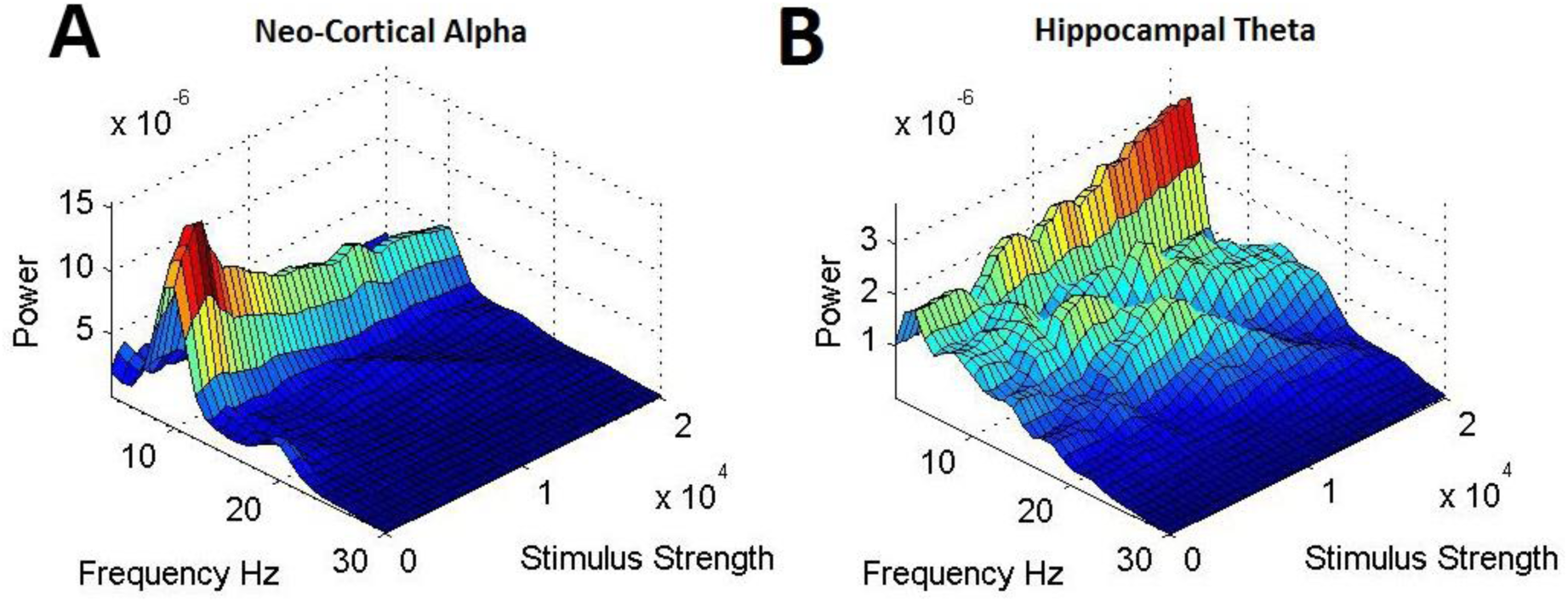
Increasing stimulus strength plotted against Neo-Cortical (NC) Alpha power **(A)** and Hippocampal Theta **(B)** power. NC Alpha power initially increases when stimulus strength has a small increase, before de-synchronising with increasing stimulus input. Hippocampal Theta power gradually increases with stimulus strength.

In contrast, the Hippocampus exhibits a strong synchronisation of 4Hz (Figure 7B) with increasing stimulus strength. This is due to the ADP function preventing neurons recovering quickly after a spike event. Not only does this mean that there will not be enough spikes during the inhibitory phase of Theta to cause de-synchronisation, but the additional spikes over the excitatory phase will cause an increase in Theta power as activity increases.

### Synch/De-Synch Predicts Learning

Having demonstrated that our model mimics the described behaviour of Alpha power decreases in the NC, and Theta power increases and phase dynamics in the Hippocampus, we now link these contrasting synchronisation behaviours with learning (see Figure 8). By varying the learning rate of STDP weight change (ε) between 0-1, it was possible to assess how the model behaves with different learning outcomes. The average of all bi-directional Hippocampal weights between subgroups P & NP are used to assess learning, i.e. the stronger the weight change the better the memory. We then calculate recall power (P response to NP + NP response to P) as a percent change in power at a particular frequency from before learning to after learning, effectively allowing us to isolate the effect of learning on power. Recall powers were binned according to the P<->NP weights of each simulation, where the distribution of weights over 1000 simulations can be seen in the inset histogram within Figure 8D. A bootstrap then provided the confidence intervals (solid lines) around a mean (dashed line) of recall power for each weight bin, allowing quadratic lines of best fit to be plotted through these points for pre-stimulus (black) and post-stimulus (red) periods.

**Figure 8:**
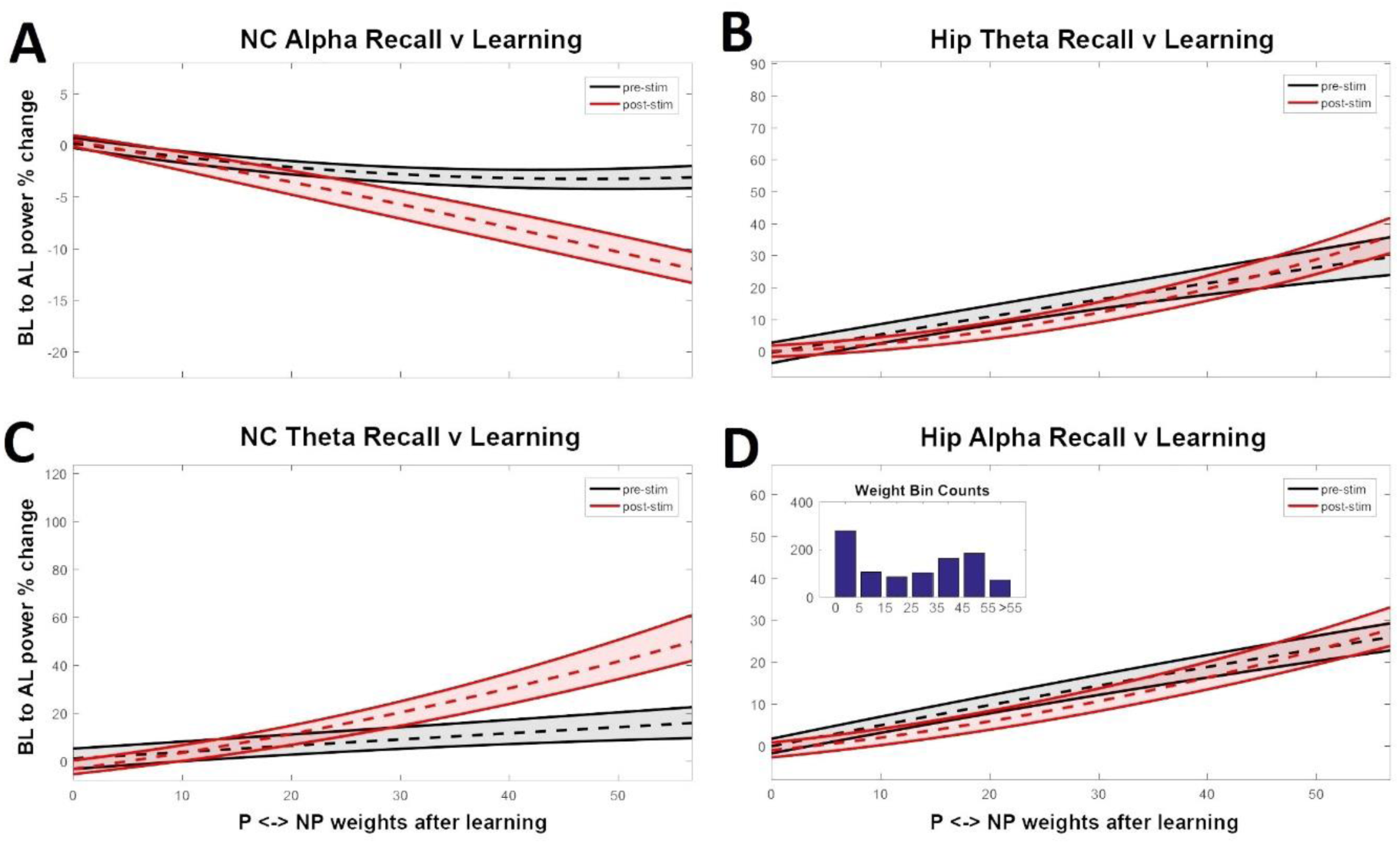
The effect of increasing synaptic efficacy between preferred (P) and non-preferred (NP) subgroups on neo-cortical (NC) alpha power **(A)**, Hippocampal Theta power **(B)**, NC Theta power **(C)** and Hippocampal Alpha power **(D)**. The inset in **D** shows the distribution of weights from 1000 simulations, where the learning rate (*ε*) was incremented gradually from 0 to 1.

From this we can use power at a particular frequency to predict whether learning has successfully occurred in our model. In respect of the sync/de-sync theory (Hanslmayr, et al., 2016), the model indicates that both a de-synchronisation of Alpha in NC areas (Figure 8A) and a synchronisation of Theta in Hippocampal areas (Figure 8B) during recall can predict successful memory retrieval.

Interestingly, one could also look at pre-stimulus Theta and Alpha power in the Hippocampus to predict whether learning has occurred (Figure 8B & 8D; black), where both increase up to 30% due to stronger weights within the Hippocampus and reciprocal connectivity between the Hippocampus and NC. This is consistent with evidence that reports the importance of pre-stimulus Theta for learning (Gyderian, et al., 2009) (Fell, et al., 2011). The effect of feedback activity plays a smaller role in NC areas, where a small decrease (<5%) in pre-stimulus Alpha power (Figure 8A; black) and an increase (<20%) in pre-stimulus Theta power (Figure 8C; black) can also predict learning. Importantly, there is a large synchronisation of Theta (<60%) at recall (Figure 8C; red) in NC areas, agreeing with experimental findings (Burke, et al., 2014) (Klimesch, et al., 2005).

## Discussion

We have presented a relatively simple spiking neural network model, which captures the complex synchronizing and desynchronizing behaviours of hippocampus and neocortex during encoding and retrieval in a typical memory task. This model, which we term the Sync/deSync (SdS) model, simulates hippocampal theta synchronization and neocortical alpha desynchronization in the service of encoding and retrieving novel stimulus associations ‒ a key requirement of episodic memory. Consistent with the notion that one-shot learning occurs in the hippocampus, but not in the neocortex (O’Reilly, et al., 2014), our model only implements synaptic modifications in the hippocampus. This hippocampal learning uses two well-described synaptic modification mechanisms. The first is spike-timing-dependent-plasticity (Song, et al., 2000), where synaptic modifications increase exponentially with decreasing time lag between the firing of pre and post-synaptic neurons. The second mechanism is theta phase-dependent plasticity, where synapses between neurons firing at the trough of theta (i.e. inhibitory phase) are strengthened, whereas synaptic connections between neurons firing at the peak of theta (i.e. excitatory phase) are weakened (Hasselmo, 2005). In the model neo-cortex, neurons fire phase-locked to an alpha oscillation at 10 Hz when they receive no input (Jensen & Mazaheri, 2010) (Klimesch, et al., 2007). When these neurons are driven by a stimulus, they increase their firing rate and gradually desynchronize from the ongoing alpha, especially when the input is strong enough to overcome maximum inhibition. Therefore, alpha power decrease is negatively related to the neural firing rate (apart from the subtle power increase at very low stimulus intensities), thereby mimicking the well-known negative relationship between alpha and neural firing (Haegens, et al., 2011).

The Sync/deSync model draws inspiration from and resonates with a number of previous models that seek to incorporate oscillations into the complementary learning systems framework. In particular, the concept of theta phase-dependent plasticity in the Hippocampus has inspired aspects of a number of influential neural models (Hasselmo, et al., 2002) (Ketz, et al., 2013) (Norman, et al., 2005). For example, Norman et al. (2005) present an important refinement of the basic complementary learning systems model, in which the strength of k Winner-Take-All (kWTA) inhibition is varied across theta phases. The effect of this modulation of inhibition is to provide a theta-phase dependent learning, with parallels to the Sync/deSync model. That is, in the Norman et al. (2005) model, the high inhibition phase of theta generates selective activation, restricting above-threshold activation to strongly responding units. LTP is then applied just to the active units, enabling selective weight update. This has similarities to the Sync/deSync idea that strongly active units move their spiking forward in the phase of theta, enabling LTP (which only obtains in the inhibitory phase) to be selectively applied.

The match between the Norman et al and Sync/deSync models for the low inhibition phase of theta is a little weaker than for the high inhibition phase, but there are still parallels. Specifically, both models exhibit activation of a broader profile of units in the low inhibition phase. In the Norman et al model, this enables LTD to be applied to competitor units (that are not strongly tuned to the memory being encoded). Sync/deSync similarly applies LTD in this low inhibition phase, however, it is a non-specifically applied, passive, decay.

Our model also takes inspiration from the influential Jensen & Lisman (2005) model, which uses phase precession to realise a working memory buffer. Our use of an ADP function to reduce the capacity for units to spike multiple times in quick succession is inherited from the Jensen & Lisman model. Additionally, while advancing the phase of theta at which a unit spikes plays a key role in the Sync/deSync model, it is somewhat different to precession in the Jensen & Lisman model, where it encodes serial order.

The Sync/deSync model is also able to capture a number of human electrophysiological findings during memory tasks. We consider these findings, starting with studies focusing on the human MTL (i.e. hippocampal layer in our model). Human single neuron recordings revealed that hippocampal neurons can change their tuning, by showing an increase in firing rate to a non-preferred stimulus after this stimulus has been associated with a preferred stimulus (Ison, et al., 2015). Furthermore, the response of these hippocampal neurons is transiently phase-locked to theta (Rey, et al., 2014). Both of these results are consistent with our modelled responses of hippocampal neurons (see Figure 3). The transient theta phase-locked response of these hippocampal neurons is due to the ADP function (Jensen, et al., 1996), which resets when a stimulus is presented, putting it out of synchrony with the ongoing theta rhythm (i.e. the 4Hz sine wave). This phase-locked response vanishes as the ADP function resynchronises to the ongoing theta, which randomly varies in phase with respect to stimulus onset and therefore averages out over trial repetitions. Such interactions between an ongoing rhythm (i.e. 4Hz sine wave) and in-built timing mechanisms (e.g. ADP function) in neurons provide a possible mechanism for the generation of transient phase-locked responses, i.e. ERPs (Mazaheri & Jensen, 2005).

Rutishauser et al. (2010) showed that a significant portion of neurons in the MTL are phase-locked to the ongoing theta rhythm during memory encoding, with an increase in theta phase-locking predicting later memory performance. Our model is consistent with these findings in showing theta phase-locked neural responses, and increased theta synchronicity to be related to later memory performance. However, Sync/deSync also suggests that responsive neurons during learning are less locked to the ongoing theta phase (Figure 4A), which seems at odds with Rutishauser et al. (2010). This decrease in theta phase-locking is present for responsive neurons only, occurring since these units overcome maximum inhibition and thus fire at the LTP phase of theta. Importantly, Rutishauser et al. (2010) did not separate neurons into stimulus responsive (i.e. showing an increase in firing rate) or not, therefore these findings cannot be directly linked to our model. However, an interesting prediction that arises from the model is that the preferred phase of firing differs between responsive and non-responsive neurons, and that this phase difference is related to later memory performance. Indeed, Rutishauser et al. (2010) found that different neurons were locked to different phases of ongoing theta. In our model, this difference is most prominent when only the first spike occurring after maximum inhibition is considered, a specific prediction that can be tested in future experiments.

Non-invasive studies in humans using MEG and source modelling showed theta power increases during successful encoding of stimulus associations in the MTL (Kaplan, et al., 2012) (Staudigl & Hanslmayr, 2013) (Backus, et al., 2016). However, there are also other studies that found the opposite pattern, i.e. MTL theta power decreases predicting successful encoding of memories (Fellner, et al., 2016) (Crespo-Garcia, et al., 2016) (Greenberg, et al., 2015). The reasons for these discrepant findings may lie in the specifics of the paradigm (i.e. type of stimuli used) or analysis (i.e. whether 1/f correction was applied). Indeed, importantly, Lega et al. (2012) found both increases and decreases in MTL theta power during successful memory encoding, with the increases occurring at a slower theta frequency (3-4Hz) and the decreases at a higher frequency (^~^8Hz). This finding is in line with MEG studies showing increased power at lower theta (3-5Hz) predicting associative memory formation (Kaplan, et al., 2012) (Staudigl & Hanslmayr, 2013) (Backus, et al., 2016). It was these studies that led us to choose a 4Hz sine wave as the pacemaker for hippocampal neurons. Together, our model nicely captures this positive relationship between 4Hz theta power and memory performance (Figure 6-8).

With respect to alpha, many studies have shown that a decrease in alpha power coincides with successful encoding and retrieval of episodic memories (see Hanslmayr et al., 2012; Hanslmayr & Staudigl, 2014 for reviews). In most previous studies, these effects extend also to beta. For this reason, and to ensure model simplicity, we have assumed only one cortical alpha rhythm, we, though, see no reason why the same principles would not also apply to beta. During successful encoding of episodic memories, alpha/beta power decreases have been found in left frontal areas for verbal material (Hanslmayr, et al., 2009) (Hanslmayr, et al., 2011) (Meeuwissen, et al., 2011) and occipital for visual material (Noh, et al., 2014). During retrieval, alpha/beta power decreases indicate the areas that are being reactivated, i.e. house the memory representation (Waldhauser, et al., 2016) (Michelmann, et al., 2016) (Khader & Rosler, 2011). This targeted alpha/beta power decrease is exactly what is modelled here, with only neural assemblies that actively process the stimulus during encoding or retrieval showing power decreases, and the degree of this power decrease predicting memory performance. An interesting model prediction is that the degree of alpha power decrease should correlate with the degree of hippocampal theta power increase, and the degree of phase precession of responsive neurons in the hippocampus. This prediction can be tested in intracranial EEG studies, which often record simultaneously from the neocortex and the hippocampus.

## Acknowledgements

This research was funded by the ERC grant Code4Memory (Grant Agreement 647954) awarded to SH.

